# Preparatory beta activity reflects the precision demands of an upcoming movement

**DOI:** 10.64898/2026.11.20.765987

**Authors:** Dominique Delisle-Godin, Pierre-Michel Bernier

## Abstract

Noise and external disturbances continuously push the limb away from its intended path, but not every deviation compromises the goal of an action. The accuracy constraints of a task define the range of acceptable endpoints and thereby set its precision demands, that is, how much noise must be attenuated for the task to succeed. Whether neural activity during movement preparation reflects these precision demands remains largely unexplored. We recorded electroencephalography (EEG) while healthy human participants (n = 27) prepared reaching movements toward narrow or wide targets at the same location and with the same timing constraints. An instructed delay before the go cue allowed preparatory activity to be isolated, and mechanical perturbations were applied during movement to probe feedback responses. Narrow targets led to more precise movements, with greater endpoint accuracy, lower variability and smaller perturbation-induced deviations. This was achieved without a reduction in movement speed, and involved greater muscle co-activation and larger long-latency responses to perturbations. Before movement onset, beta power (13-30 Hz) was lower for narrow than for wide targets, an effect that emerged late in the delay period and was concentrated over bilateral parieto-occipital electrodes. In contrast, alpha (8-12 Hz) and theta (4-7 Hz) activity did not differ between conditions. These findings demonstrate that preparatory beta activity is sensitive not only to kinematic requirements of the movement but also the precision with which it must be produced.

**Significance Statement:** Before a movement begins, the nervous system prepares where and how fast to move, but whether it also prepares how precisely to move remains unclear. We show that narrowing the target of a reaching movement, while keeping target location and timing constraints the same, leads to more precise movements, with noise attenuated through greater muscle co-activation and stronger feedback responses. Most importantly, narrow targets lowered preparatory beta-band activity before movement onset, whereas alpha and theta activity were unaffected. Preparatory beta is known to be modulated when movement parameters such as speed must change. These findings show that it is also sensitive to precision demands, suggesting that movement preparation takes into account how precisely a movement must be executed.

## Introduction

Movements are corrupted by noise arising from the execution of motor commands, in the neural processes that plan them, and from external disturbances (Churchland et al., 2006b; Faisal et al., 2008). Yet which deviations matter depends on the accuracy constraints of the task, as lateral deviations must be attenuated with a narrow target but are inconsequential with a wide one. Because attenuating noise is costly (Manohar et al., 2015), movements are only as precise as required. With a wide target, endpoint variability is larger and perturbation-induced deviations are corrected less than with a narrow target (Liu and Todorov, 2007; Knill et al., 2011), as formalized by the minimal intervention principle of optimal feedback control (Todorov and Jordan, 2002). Precision is thus actively controlled through several mechanisms (Codol et al., 2020). Co-activation of antagonist muscles stiffens the limb, attenuating the mechanical effects of noise and perturbations (Gribble et al., 2003). Feedback control corrects deviations using an estimate of the limb’s state that combines sensory feedback with efference-copy predictions (Wolpert et al., 1995; Todorov and Jordan, 2002). A higher feedback gain strengthens corrections of deviations detected through sensory feedback, expressed as early as the long-latency response (LLR) to perturbations (Pruszynski and Scott, 2012; Nashed et al., 2012), and of deviations caused by noise in the motor command, detected through internal feedback (Chen-Harris et al., 2008; Manohar et al., 2019).

Movements are prepared before they are executed (Tanji and Evarts, 1976; Churchland and Shenoy, 2024), and their precision may be specified during preparation. First, variability in preparatory activity predicts variability of the upcoming movement (Churchland et al., 2006b), suggesting that precision is constrained in part by preparation. Second, feedback gains are already tuned to task demands before movement onset (Papaioannou and Dimitriou, 2021; Torell et al., 2023; Rohlén et al., 2025). Preparatory activity could therefore reflect not only the parameters of the upcoming movement but also its required precision. Yet studies of preparatory activity have mostly contrasted movements differing in direction (Tanji and Evarts, 1976; Crammond and Kalaska, 2000), extent (Messier and Kalaska, 2000), force (Riehle et al., 1994) or speed (Churchland et al., 2006a). Target size has also been varied, but with movement speed free to vary (Gomez et al., 2000), leaving precision demands not fully isolated. Whether preparatory activity differs as a function of precision demands when target location and timing constraints prevent simply slowing down therefore remains unclear.

EEG is well suited to track activity during the delay preceding a movement, and beta-band (13-30 Hz) event-related desynchronization (ERD) over sensorimotor regions is a classic marker of movement preparation (Pfurtscheller and Lopes da Silva, 1999; Kilavik et al., 2013). Its magnitude varies with movement speed (Pierrieau et al., 2023, 2025), adaptation to perturbations (Torrecillos et al., 2015; Alayrangues et al., 2019; Jahani et al., 2020), directional uncertainty (Tzagarakis et al., 2010, 2015) and reward (Savoie et al., 2019). Notably, performance-contingent rewards lead to more precise movements (Manohar et al., 2015; Savoie et al., 2019; Codol et al., 2020), and lower preparatory beta power (Savoie et al., 2019), raising the possibility that preparatory beta is sensitive to precision demands, although this has not been directly tested. Higher precision demands could also engage more general preparatory processes, such as greater visuospatial attention, associated with posterior alpha (8-12 Hz) desynchronization (Jensen and Mazaheri, 2010; Klimesch, 2012), or greater cognitive control, associated with midfrontal theta (4-7 Hz) activity (Cavanagh and Frank, 2014).

Here, we asked whether preparatory activity reflects precision demands, manipulated via target width. Participants reached toward narrow or wide targets at the same location and with the same timing constraints, while EEG was recorded during an instructed delay. Endpoint accuracy and variability assessed movement precision, and co-activation and LLRs to perturbations assessed the mechanisms attenuating noise. We predicted lower preparatory beta for narrow than for wide targets, and examined alpha and theta to test whether any effect was specific to beta.

## Materials and Methods

### Participants

Twenty-seven healthy, right-handed adults (13 females; age 19-32 years) participated in the study. All participants reported no history of musculoskeletal or neurological disorders and had normal or corrected-to-normal vision. Experimental procedures were approved by the Ethics Committee of the Université de Sherbrooke, and all participants provided written informed consent. Participants received monetary compensation (25$ CAD plus a performance-based bonus of 5-10$ CAD). Total session duration was approximately 1.5 h.

### Apparatus

Participants performed planar reaching movements using a KINARM End-Point Lab (BKIN Technologies, Kingston, ON). While seated, participants held the robotic manipulandum handle and controlled a cursor representing their hand. Direct vision of the arm was occluded. This setup allowed controlled mechanical loads to be applied to the handle and precise control of the visual feedback.

### Experimental procedure

Fig. 1A,B illustrates the workspace and trial timeline. Each trial began with the participant moving the cursor into a start position (target radius 1 cm) located 5 cm from the proximal edge of the workspace. Participants fixated a central cross and maintained position for 1.5 s to establish baseline EEG activity. A reach target located 20 cm away from the start position then appeared, and participants remained at the start position until the go signal, indicated by the target border turning green after a 1 s delay. Participants were instructed to reach the target with a movement time of 200-400 ms. No reaction time constraints were imposed. Once the target was reached, participants were required to maintain the cursor within the target for 250 ms, after which feedback was displayed. The target turned green for accurate reaches and red if participants failed to stabilize the cursor in the target. Movement time feedback (“too fast”, “too slow” or “good speed”) was also provided. Participants earned 10 points for trials that were both accurate and within the desired movement-time window.

**Figure 1.**
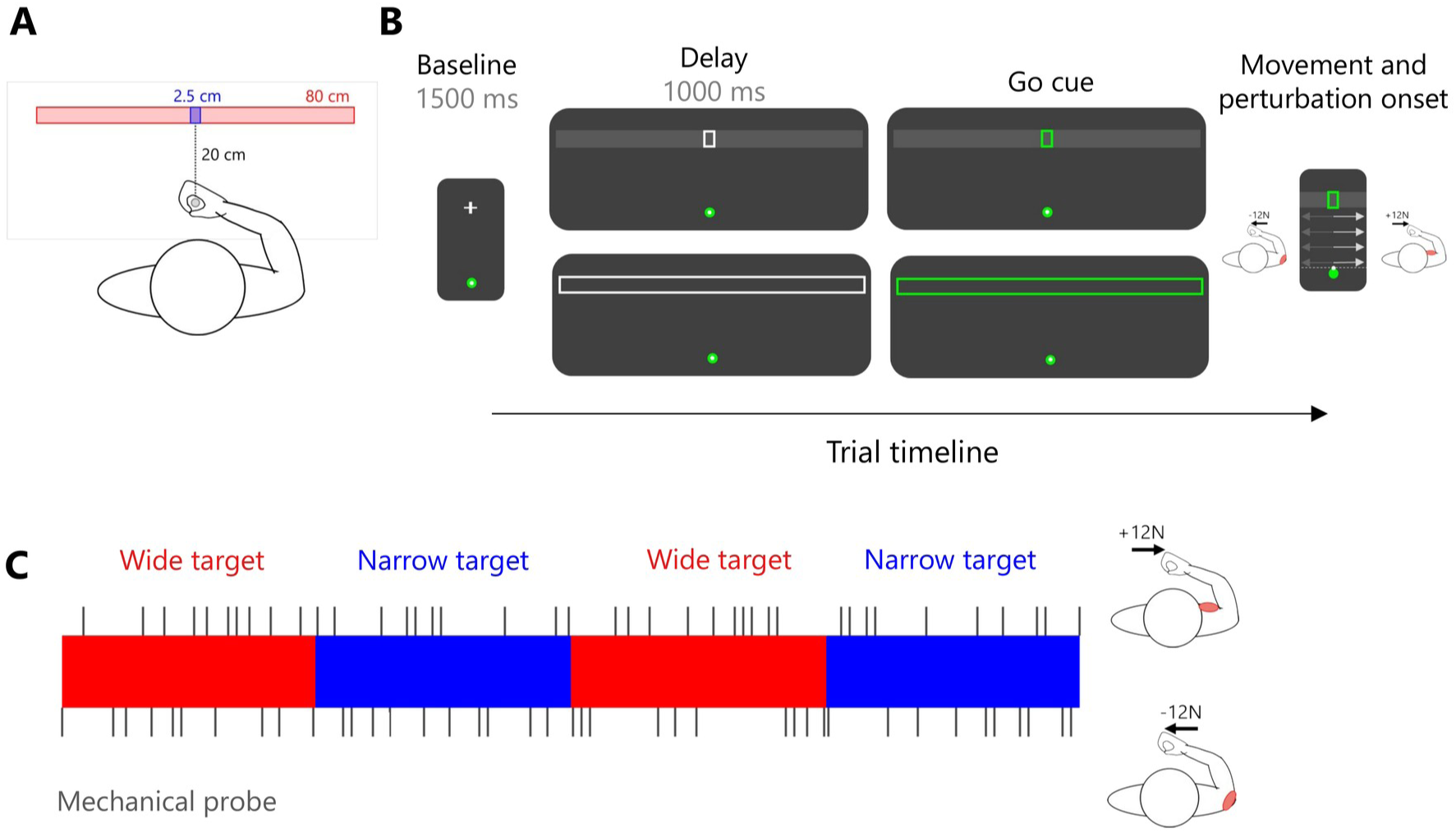
Experimental setup and design. A, Participants made reaching movements with a robotic manipulandum from a start position to a target located 20 cm away. The target was either narrow (2.5 cm) or wide (80 cm). In the narrow target condition, the narrow target was displayed over a grey wide target. B, Trial timeline. After a 1500 ms baseline with central fixation, the narrow (top) or wide (bottom) target appeared, and participants waited through a 1000-ms delay until the target turned green (go cue). Mechanical perturbations were applied on probe trials shortly after movement onset. C, Schematic of the blocked design. Vertical lines above and below each block indicate probe trials with rightward (+12 N) and leftward (-12 N) loads. Insets show the direction of each load.

### Main experimental conditions

Target width was varied across two conditions, imposing tighter accuracy constraints with a narrow target and looser accuracy constraints with a wide target. In the wide-target condition, the target was 80 cm wide (240 trials), whereas in the narrow-target condition it was 2.5 cm wide (240 trials). Both targets were 4 cm in height. To minimize differences in the visual display between conditions, a grey wide target was displayed underneath the narrow target in the narrow-target condition.

Half of the trials (240 total, 120 per target condition) were performed with online visual feedback of the cursor during movement, whereas the other half was performed without visual feedback. Vision and no-vision conditions were conducted in separate blocks. Results from the no-vision condition were qualitatively similar to those obtained with visual feedback. To maintain focus on the primary hypotheses, the main text reports analyses from trials performed with visual feedback, whereas key results from the no-vision condition are presented in Extended Data Figure 4-2.

Corrective responses were probed by applying mechanical loads (step perturbations; 10 ms linear ramp-up) to the manipulandum when participants reached a predetermined location just outside the start position (Fig. 1B). A +12 N load displaced the hand rightward, and a -12 N load displaced it leftward. Perturbations were delivered pseudo-randomly within each block, with 20 probe trials (10 per direction). Probes were unpredictable and never occurred on consecutive trials, yielding 40 probes per target-width condition.

The experiment used a blocked design in which participants completed blocks of 60 trials with one target size and alternated between wide and narrow target blocks. The order of the blocks (wide first vs narrow first) was pseudo-randomized and counterbalanced across participants.

### Kinematic data recording and analysis

Kinematic data were recorded continuously using the KINARM system at 1000 Hz and low-pass filtered using a second-order, dual-pass Butterworth filter with a 30 Hz cutoff. Movement onset was defined as the time at which the cursor exited the start position, and movement end as the time at which the cursor had remained within the target for at least 10 ms. Reaction time was defined as the interval between the go cue and movement onset, and movement time as the interval between movement onset and movement end. Trials with reaction times longer than 1 s were excluded (72 trials, 0.56% of total). For unperturbed trials, movement speed was characterized by peak forward velocity and time to peak velocity relative to movement onset. Endpoint accuracy was quantified as endpoint error, defined as the absolute x-axis distance from the target center at movement end. Lateral variability was defined as the standard deviation of lateral position across trials and was computed both at peak velocity and at movement end (endpoint variability). On probe trials, perturbation-induced deviations were quantified as the maximum lateral deviation relative to the mean lateral trajectory of unperturbed trials (Δx max) and the change in lateral endpoint position relative to the mean endpoint of unperturbed trials (Δ endpoint).

### EMG data recording and analysis

Surface electromyography (EMG) was recorded from the pectoralis major and posterior deltoid using a Delsys Trigno wireless system (Delsys Inc., Boston, MA). These muscles were selected because they are robustly recruited by similar perturbations (Crevecoeur et al., 2019; De Comite et al., 2022). Before electrode placement, the skin was shaved and cleaned with isopropyl alcohol, and electrodes were positioned over the muscle belly and aligned with fiber orientation. EMG signals (±5 V output range, effective gain ≈ 909 V/V) were sampled at 1000 Hz, synchronized with kinematic data, and band-pass filtered online (20-450 Hz). Offline, signals were band-pass filtered with a third-order, zero-lag Butterworth filter (20-450 Hz) and full-wave rectified. EMG was normalized to activity recorded during a calibration procedure performed at the beginning of the experiment, in which participants maintained the start position against a +12 N (pectoralis major) or -12 N (posterior deltoid) load for 10 s, with five trials per load. In unperturbed trials, co-activation of the two muscles was quantified around movement onset (Crevecoeur et al., 2019). At each time point, co-activation was defined as the smaller of the two normalized EMG signals, which corresponds to the level of activity shared by both muscles. This value was then averaged over a 200-ms window centered on movement onset. In perturbed trials, perturbation-evoked muscle responses were quantified by subtracting the mean EMG of unperturbed trials from the EMG of perturbed trials (ΔEMG). ΔEMG was averaged within short-latency (20-50 ms), long-latency (50-105 ms) and voluntary (120-180 ms) windows after perturbation onset (Pruszynski et al., 2008), with LLRs used as the primary readout of feedback gains (Pruszynski and Scott, 2012). Responses were analyzed in the agonist muscle for each perturbation direction, namely the posterior deltoid for -12 N loads and the pectoralis major for +12 N loads.

### EEG data recording and analysis

Scalp EEG was recorded using a 64-channel BrainVision actiCAP system (extended 10-20 layout; Brain Products) and a BrainAmp amplifier at 500 Hz. Preprocessing was performed in MATLAB using EEGLAB (Delorme and Makeig, 2004). Data were band-pass filtered between 1 and 49 Hz, and noisy channels were removed. The data were down-sampled to 128 Hz and epoched from -2.5 to +1.5 s relative to the go cue. Independent component analysis (runica) was used to identify and remove ocular and muscle artifacts. After ICA, data were re-referenced to the average of all electrodes, and removed channels were interpolated.

Because target width differed between conditions and could therefore produce condition-dependent visually evoked activity following target presentation, we removed the event-related potential (ERP) from single trials before time-frequency decomposition. This step allowed us to focus on non-phase-locked (induced) oscillatory activity and minimized the contribution of differences in evoked responses to subsequent comparisons of oscillatory power. For each participant and condition, the ERP was computed as the average of all retained trials of that condition (perturbed and unperturbed trials pooled), and this condition-specific ERP was subtracted from each of those trials (Cohen, 2014).

Time-frequency analysis was then performed using FieldTrip (Oostenveld et al., 2011) with Morlet wavelets (3-10 cycles) from 4 to 44 Hz in 1-Hz steps. Power was averaged by condition (all perturbed and unperturbed trials per condition) and expressed as relative change from a 400 ms baseline preceding target onset (-1600 to -1200 ms).

### Experimental design and statistical analysis

All statistical analyses were conducted with custom MATLAB and Python scripts. Kinematic and EMG variables were compared between target-width conditions (wide vs narrow) using within-subject tests. Normality was assessed with the Shapiro-Wilk test. For normally distributed data, two-tailed paired t-tests were used; otherwise, Wilcoxon signed-rank tests were applied. The significance level was set at α = 0.05. Exact p values are reported unless p < 0.001. Effect sizes are reported as Cohen’s d for paired samples.

Condition-related differences in oscillatory activity were assessed with cluster-based permutation tests implemented in FieldTrip (Maris and Oostenveld, 2007; Oostenveld et al., 2011). We focused on theta (4-7 Hz), alpha (8-12 Hz) and beta (13-30 Hz) bands and contrasted time-frequency power between conditions in the -1 to 0 s interval preceding the go cue. Beta was the band of primary interest, based on the a priori hypothesis that preparatory beta power would be lower for narrow than for wide targets. Alpha and theta were analyzed as control bands to assess whether any effect was specific to beta, and each band was tested separately. Dependent-samples t-tests were computed at each time-sensor point, and samples with p < 0.05 were grouped into clusters based on spatial adjacency (each channel required ≥2 neighbors). Cluster-level statistics were defined as the sum of t-values within each cluster and evaluated against a null distribution generated by 1000 random permutations of condition labels. Clusters exceeding the 97.5th percentile of the null distribution were deemed significant (α = 0.05).

## Results

Reaction times, computed from all trials because perturbations occurred only after movement onset, were slightly but significantly shorter for the narrow target (345 ms) than for the wide target (358 ms; W = 94, *p* = 0.021, *d*_z_ = -0.39).

### Behavior during unperturbed trials

Inspection of hand trajectories (Fig. 2A) showed a small rightward bias in the wide-target condition, whereas movements in the narrow-target condition ended closer to the target center (wide 2.11 cm vs narrow 0.21 cm). Because the wide target did not require participants to counter this natural rightward tendency of the right arm, the endpoint shift can be interpreted as tolerance of a task-irrelevant bias. Reflecting this, endpoint accuracy was greater for the narrow target, with mean endpoint error differing significantly between conditions (wide 2.46 cm vs narrow 0.58 cm; W = 0, *p* < 0.001, *d*_z_ = -1.34; Fig. 2B).

**Figure 2.**
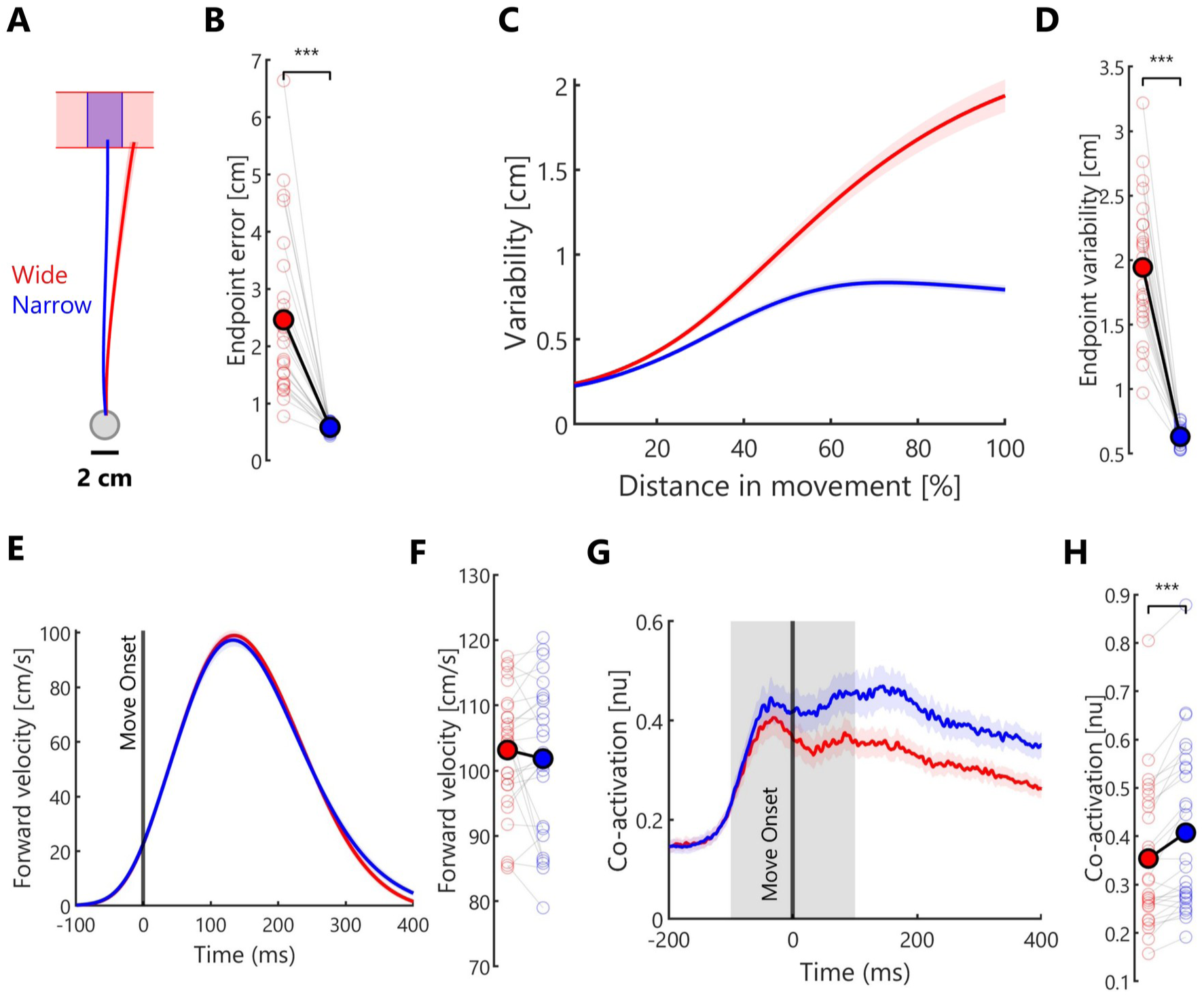
Narrow targets led to more precise movements in unperturbed trials without a reduction in movement speed. A, Mean hand trajectories for the wide (red) and narrow (blue) targets. B, Lateral endpoint error. C, Movement variability as a function of the distance traveled (percentage of movement). D, Endpoint variability (standard deviation of lateral endpoint position). E, Forward velocity aligned to movement onset. F, Peak forward velocity. G, Co-activation aligned to movement onset. The grey area indicates the 200 ms window centered on movement onset. H, Co-activation averaged over this window. In time courses, lines show the group mean and shaded areas the SEM. In individual-data panels, light markers show individual participants and dark markers the group mean. n = 27. ***p < 0.001.

Examining the standard deviation of lateral hand position across trials (Fig. 2C) revealed that lateral variability increased early in the movement in both conditions (Fig. 2C). It then decreased toward movement end in the narrow-target condition, whereas it continued to increase in the wide-target condition. In line with the minimal intervention principle, endpoint variability was substantially greater for the wide target (1.94 cm) than for the narrow target (0.63 cm; *t_(26)_* = - 13.74, *p* < 0.001, *d*_z_ = -2.64; Fig. 2D). This difference was already present early in the movement, as lateral variability at peak velocity was significantly greater for the wide target (1.11 cm) than for the narrow target (0.73 cm; *t_(26)_* = -8.67, *p* < 0.001, *d*_z_ = -1.67). This suggests that movements were already less precise early in the movement with the wide target.

Greater precision with the narrow target was not achieved by reducing movement speed. Peak forward velocity did not differ between conditions (narrow 101.83 cm/s vs wide 103.15 cm/s; *t_(26)_* = -0.88, *p* = 0.385, *d*_z_ = -0.17; Fig. 2F), nor did time to peak velocity (narrow 144 ms vs wide 144 ms; W = 160, *p* = 0.500, *d*_z_ = 0.06). Movement time was significantly longer for the narrow target (294 ms) than for the wide target (276 ms; W = 39, *p* < 0.001, *d*_z_ = 0.78). This difference likely reflects the additional time required to home in on the narrow target, with more time devoted to online corrections when accuracy constraints were tighter. In both conditions, movement times remained well within the 200-400 ms window imposed by the task.

Co-activation around movement onset was significantly greater for the narrow target (0.41 normalized unit; nu.) than for the wide target (0.35 nu.; *t_(26)_* = 5.64, *p* < 0.001, *d*_z_ = 1.09; Fig. 2H). Because co-activation of antagonist muscles increases limb stiffness, this indicates that participants stiffened the limb, attenuating the mechanical effects of noise and perturbations, when accuracy constraints were tighter, a strategy known to improve movement accuracy (Gribble et al., 2003).

### Behavior during perturbed trials

Hand trajectories following perturbations (Fig. 3A) showed that participants largely tolerated perturbation-induced deviations in the wide-target condition and did not recenter the hand, whereas deviations were actively corrected in the narrow-target condition. This was reflected in the change in lateral endpoint position relative to unperturbed trials (Δ endpoint; Fig. 3B), which was significantly larger for the wide target (*t_(26)_* = -21.58, *p* < 0.001, *d*_z_ = -4.15). On average, perturbations shifted the endpoint by 10.70 cm in the wide-target condition but by only 0.92 cm in the narrow-target condition.

**Figure 3.**
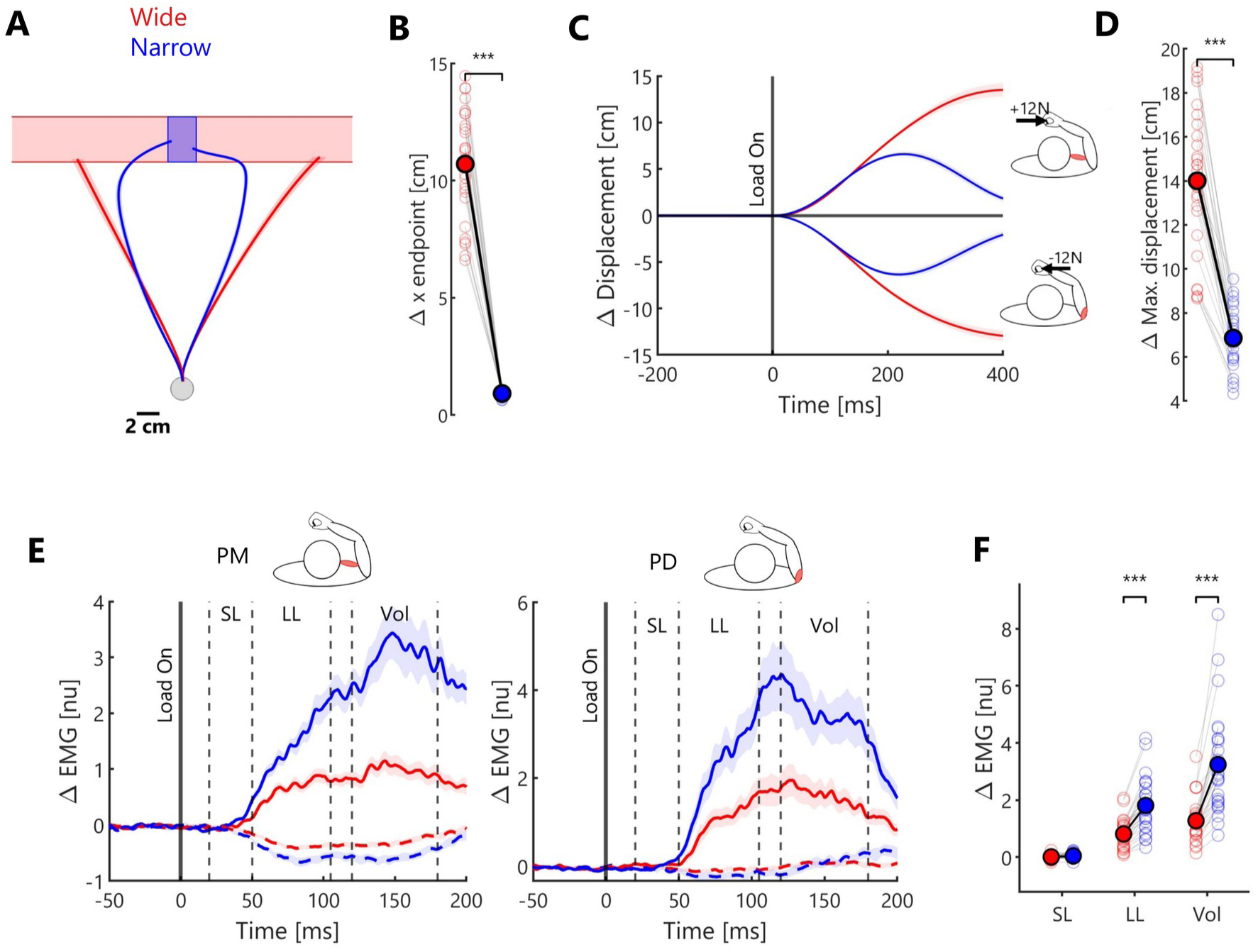
Narrow targets led to stronger corrections of perturbation-induced deviations. A, Mean hand trajectories on perturbed trials for the wide (red) and narrow (blue) targets. B, Change in lateral endpoint position relative to unperturbed trials (Δ endpoint). C, Lateral displacement relative to unperturbed trials, aligned to load onset, for rightward (+12 N) and leftward (-12 N) loads. D, Maximum lateral deviation (Δx max). E, Perturbation-evoked muscle responses (ΔEMG) in pectoralis major (PM) and posterior deltoid (PD), aligned to load onset. Solid lines show responses to the load that stretched the muscle and dashed lines responses to the opposite load. Vertical dashed lines delimit the short-latency (SL, 20-50 ms), long-latency (LL, 50-105 ms) and voluntary (Vol, 120-180 ms) windows. F, ΔEMG of the agonist muscle averaged within each window. Lines and shaded areas show the group mean ± SEM. Light markers show individual participants and dark markers the group mean. n = 27. ***p < 0.001.

A similar pattern was observed for maximum lateral deviation (Δx max; Fig. 3C,D). The perturbation displaced the hand substantially more in the wide-target condition (14 cm) than in the narrow-target condition (6.86 cm; *t_(26)_* = -14.43, *p* < 0.001, *d*_z_ = -2.78), consistent with greater attenuation of perturbation-induced deviations when accuracy constraints were tighter.

Fig. 3E shows the time course of perturbation-evoked muscle responses, aligned to load onset, with the short-latency, long-latency and voluntary windows indicated. Short-latency responses did not differ between conditions (*t_(26)_* = 1.73, *p* = 0.096, *d*_z_ = 0.33). In contrast, LLRs and voluntary responses were significantly larger in the narrow-target condition (LLR: *t_(26)_* = 9.59, *p* < 0.001, *d*_z_ = 1.85; voluntary: W = 0, *p* < 0.001, *d*_z_ = 1.58).

Together, the unperturbed and perturbed results show that narrow targets led to more precise movements without a reduction in movement speed, with noise attenuated through greater co-activation and larger LLRs.

### Pre-movement EEG activity

Given the robust effects of accuracy constraints on behavior, we next tested whether pre-movement EEG activity differed between target conditions. Time-frequency power was computed from all trials of each condition, pooling perturbed and unperturbed trials, since perturbations were applied only after movement onset and could not affect activity during the delay. Conditions were compared with nonparametric cluster-based permutation tests (Maris and Oostenveld, 2007), over the delay period (-1 to 0 s relative to the go cue). Tests were run separately for the theta (4-7 Hz), alpha (8-12 Hz) and beta (13-30 Hz) bands, with the contrast defined as narrow minus wide. Because cluster-based tests do not establish the precise latency or spatial extent of an effect (Sassenhagen and Draschkow, 2019), the timing and topography reported below describe where effects were most pronounced.

Because conditions were blocked and power was expressed relative to a pre-target baseline (Pfurtscheller and Lopes da Silva, 1999; Cohen, 2014), differences in baseline activity between blocks could in principle produce apparent differences during the delay. We therefore first compared raw power between conditions during the 1 s preceding target onset. Cluster-based tests revealed no significant differences in any of the three bands (largest cluster *p* = 0.1). Differences identified during the delay can therefore be attributed to changes in delay-period activity rather than to differences in baseline.

Because conditions differed in target width, they also produced different visually evoked responses following target presentation (Extended Data Fig. 4-1). To focus on induced oscillatory activity rather than phase-locked visual responses, condition-specific ERPs were removed prior to time-frequency decomposition (see Methods). The analyses below therefore reflect differences in induced oscillatory power.

### Beta band (13-30 Hz)

Beta-band activity desynchronized after target onset in both conditions, with the strongest overall ERD over left centroparietal electrodes contralateral to the reaching arm (Fig. 4A), consistent with sensorimotor preparation (Pfurtscheller and Lopes da Silva, 1999; Kilavik et al., 2013). The condition contrast revealed a significant difference during the delay (*t*_sum_ = -1977.78, *p* = 0.028; Fig. 4B,C). Beta power was lower for the narrow than the wide target, with the observed difference concentrated approximately 625 to 109 ms before the go cue and distributed mainly over bilateral parieto-occipital electrodes. Thus, the target-width effect was topographically distinct from the location of maximal ERD common to both conditions. Fig. 4C illustrates the time course averaged over cluster electrodes. Because these electrodes were selected from the same contrast, the traces are descriptive and were not subjected to additional testing (Kriegeskorte et al., 2009). In both conditions, beta power decreased after target onset. The two time courses were similar early in the delay and diverged during its second half, with lower beta power in the narrow-target condition.

**Figure 4.**
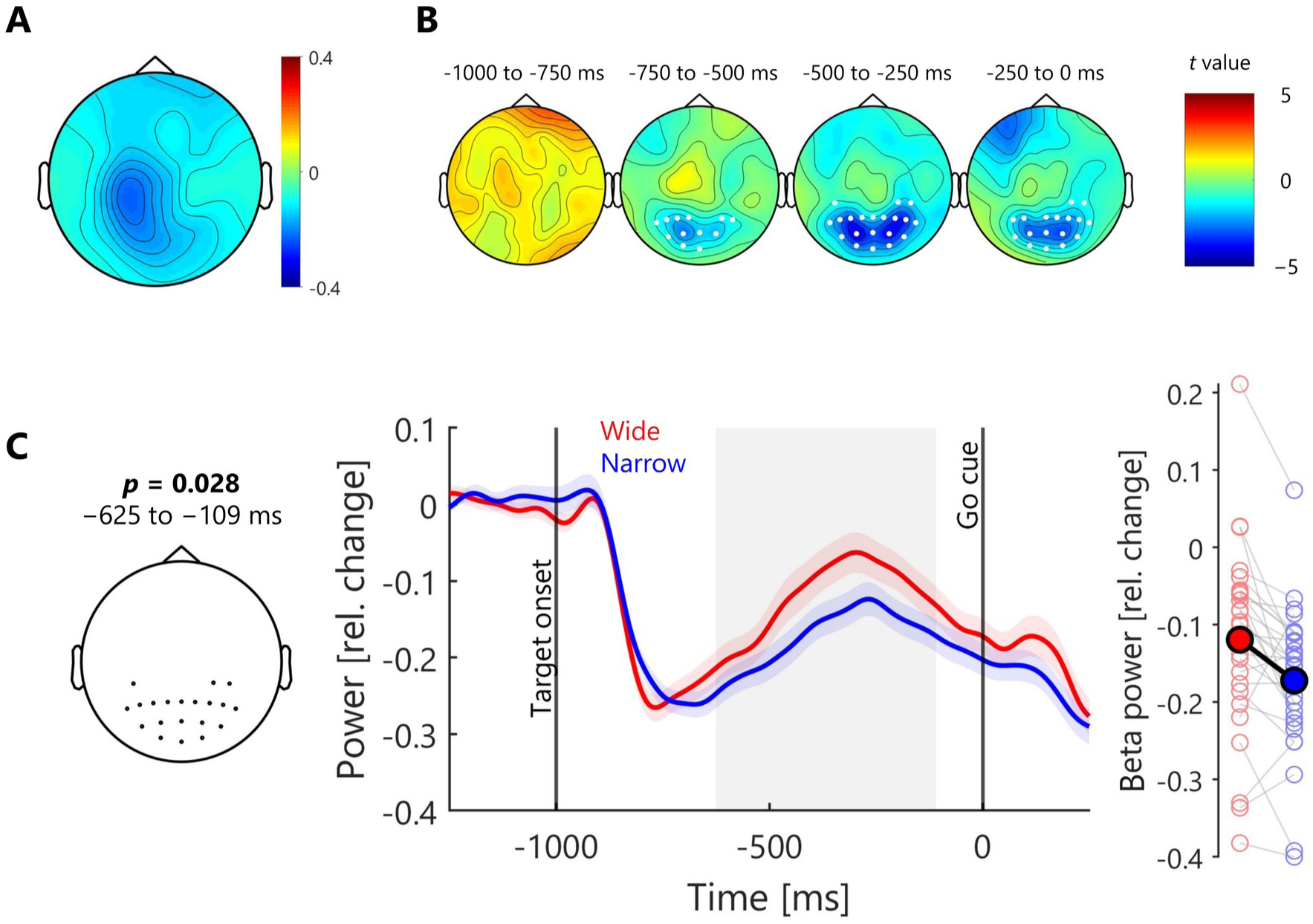
Preparatory beta power was lower for narrow than for wide targets. A, Topography of beta power (13-30 Hz, relative change from baseline) during the delay period, averaged across conditions. B, Topographies of the condition difference (narrow minus wide, t values) in four successive 250-ms windows preceding the go cue. White markers indicate electrodes belonging to the significant cluster. C, Left, electrodes of the significant cluster, with the cluster p value and time window. Middle, beta power averaged over cluster electrodes for the wide (red) and narrow (blue) targets, aligned to the go cue (mean ± SEM). The grey area indicates the time window of the cluster. Right, beta power averaged over cluster electrodes and time window for each participant (light markers) and the group mean (dark markers). Because electrodes were selected from the same contrast, the traces in C are descriptive (Kriegeskorte et al., 2009). n = 27.

Similar effects were observed in the no-vision condition (Extended Data Fig. 4-2), with significantly lower beta power for the narrow than the wide target (*t*_sum_ = -1064.03, *p* = 0.042), approximately 523 to 0 ms before the go cue and also distributed over parieto-occipital electrodes. This result not only adds robustness to the main beta finding by replicating it in an independent dataset, but it also suggests that the modulation of preparatory beta activity by target width is unlikely to be contingent on the anticipation of visually guided corrections.

### Alpha band (8-12 Hz)

Target presentation was followed by alpha-band desynchronization, mainly over parieto-occipital electrodes (Fig. 5A). No condition difference survived cluster correction (Fig. 5B). The largest, nonsignificant cluster (*t*_sum_ = 1902.6, *p* = 0.084) occurred approximately 921 to 465 ms before the go cue and extended across frontal, parietal, and occipital electrodes. Its positive sign reflects lower alpha power in the wide-target condition relative to the narrow-target condition (Fig. 5C). Thus, any tendency toward a condition effect was in the direction of greater alpha desynchronization for the wide target. Because the effect did not reach significance, these observations are descriptive only.

**Figure 5.**
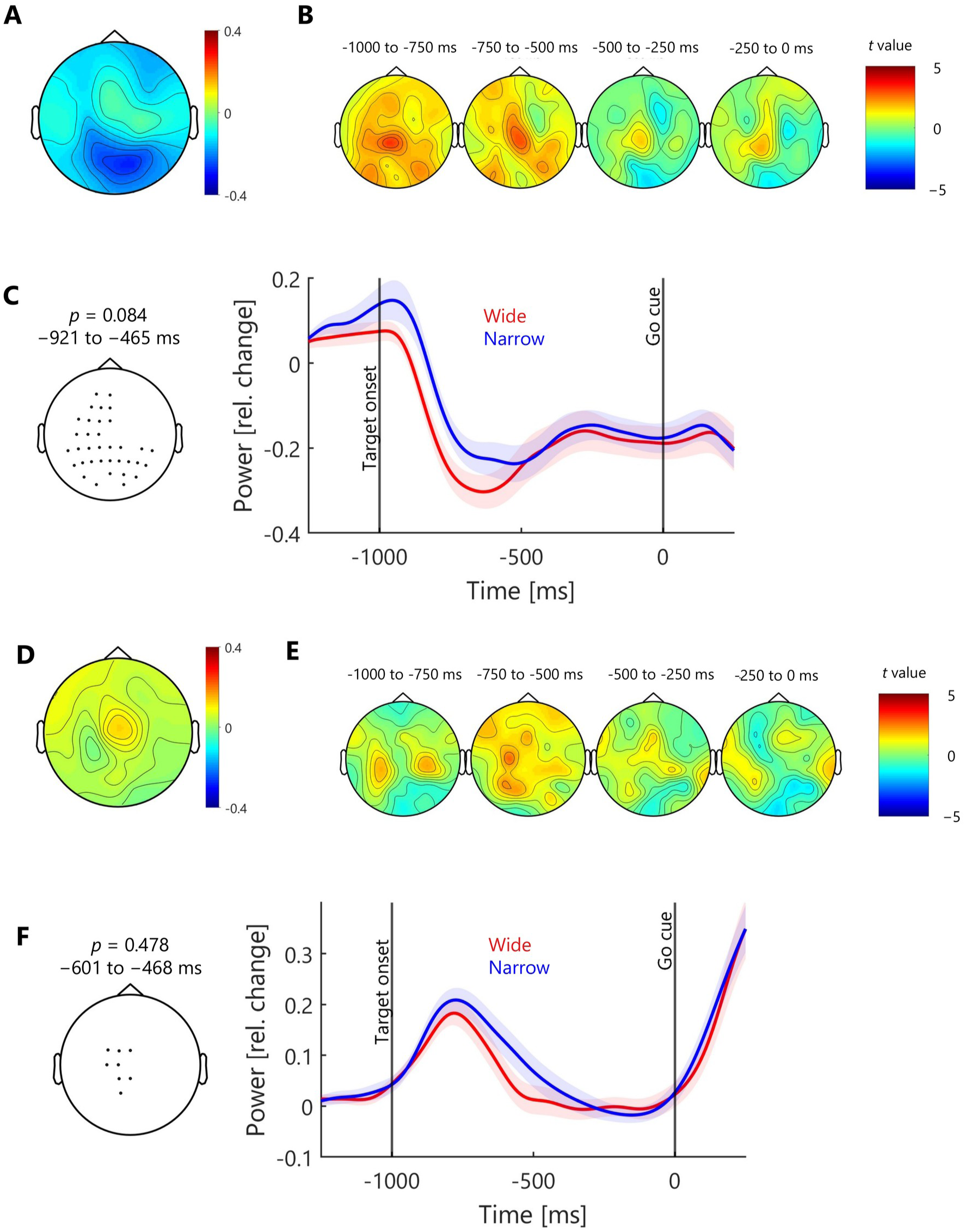
Preparatory alpha and theta power did not differ reliably between conditions. A, Topography of alpha power (8-12 Hz, relative change from baseline) during the delay period, averaged across conditions. B, Topographies of the condition difference in alpha power (narrow minus wide, t values) in four successive 250 ms windows preceding the go cue. C, Left, electrodes of the largest, nonsignificant alpha cluster, with its p value and time window. Right, alpha power averaged over these electrodes for the wide (red) and narrow (blue) targets, aligned to the go cue (mean ± SEM). D-F, Same as A-C for theta power (4-7 Hz). n = 27.

### Theta band (4-7 Hz)

Theta-band activity increased over midfrontal electrodes after target onset in both conditions (Fig. 5D). No condition difference survived cluster correction (Fig. 5E). The largest cluster was small and nonsignificant (*t*_sum_ = 204.59, *p* = 0.478) and was centered over frontocentral electrodes. Inspection of the cluster trace (Fig. 5F) revealed only minor differences between conditions. The present data therefore provide no evidence that preparatory theta activity differed as a function of target width.

Together, these results indicate that pre-movement EEG activity differed between accuracy constraints, with a significant difference observed only in the beta band.

## Discussion

We asked whether preparatory activity reflects precision demands, which we manipulated through target width. With a narrow target, movements were more precise, with greater endpoint accuracy and lower lateral variability, without a reduction in peak velocity. Noise was attenuated through several mechanisms, including larger LLRs and greater co-activation. Before movement onset, beta power was lower for narrow than for wide targets, whereas alpha and theta did not differ. Preparatory beta thus appears sensitive to the precision demands of the task.

### Preparatory beta reflects precision demands

Target location and timing constraints were identical across conditions, and peak velocity did not differ. The conditions differed mainly in how much deviation could be tolerated, and therefore in how much noise had to be attenuated. Preparatory beta is known to change when success requires changing movement parameters, such as speed (Pierrieau et al., 2023, 2025), or adapting to perturbations (Torrecillos et al., 2015; Alayrangues et al., 2019; Jahani et al., 2020). Here, success required neither a change in speed nor in aiming direction, but the attenuation of task-relevant variability, providing novel evidence that preparatory beta is sensitive to the required precision of the upcoming movement. This parallels the influence of performance-contingent incentives, which make movements more accurate and less variable (Manohar et al., 2015; Savoie et al., 2019; Codol et al., 2020) and lower preparatory beta (Savoie et al., 2019), an effect attributed to the motivational value of reward. As reward was held constant here, the present effect is unlikely to reflect motivational value alone. Both findings can be reconciled if lower preparatory beta accompanies a greater allocation of resources to precision, which reward makes worth its cost (Manohar et al., 2015) and a narrow target makes necessary for success. This fits evidence that premovement beta tracks the neurocomputational cost of an action rather than its speed (Pierrieau et al., 2023).

### Lower preparatory beta and the attenuation of noise

Functionally, beta activity has been proposed to favor maintenance of the current sensorimotor state, or “status quo” (Engel and Fries, 2010). Conversely, reduced beta power has been suggested to increase the computational capacity of task-relevant neuronal populations, allowing them to express more differentiated patterns of activity and to process incoming information more flexibly (Engel and Fries, 2010; Brittain et al., 2014). On this view, lower preparatory beta may accompany a greater allocation of neural resources to the upcoming action (Savoie et al., 2019; Pierrieau et al., 2023), such as those needed to attenuate intrinsic noise, which improves precision but is costly (Manohar et al., 2015). The need for these resources may drive beta desynchronization, or beta desynchronization may be required to deploy them. With a narrow target, these resources could attenuate noise in two non-exclusive ways.

First, lower beta could accompany a more precisely specified action. In dynamical systems accounts, preparation brings motor cortical activity toward an optimal subspace of states from which the movement unfolds, a process indexed by a decline in trial-to-trial variability (Churchland et al., 2006c). Residual variability in this preparatory state predicts part of the variability of the upcoming movement (Churchland et al., 2006b), and preparatory states pushed beyond the optimal region are associated with poorer performance (Smoulder et al., 2024). How precisely this state can be set depends on how fully the action can be specified. When several movements remain possible, preparatory activity settles in an intermediate state between them (Dekleva et al., 2018; Meirhaeghe et al., 2023). Correspondingly, preparatory beta desynchronization is attenuated in such conditions (Tzagarakis et al., 2010, 2015). A wide target likewise leaves many endpoint states compatible with success (Padmanabhan et al., 2026), whereas a narrow target restricts them, which might require the preparatory state to be set more precisely. Lower beta with the narrow target may thus accompany stronger quenching of preparatory variability, leaving preparation less corrupted by internal noise. Consistent with this, lateral variability was already lower at peak velocity for narrow targets.

Second, lower beta could accompany the setting of higher feedback gains. In optimal feedback control, corrections are computed from an estimate of the limb’s state that combines efference-copy predictions with sensory feedback (Wolpert et al., 1995; Todorov and Jordan, 2002), so that the same feedback gains correct deviations detected through sensory feedback, as for perturbations and peripheral noise, or through internal feedback, as for noise in the motor command (Chen-Harris et al., 2008; Manohar et al., 2019). Consistent with this, corrections were stronger with the narrow target in both perturbed and unperturbed movements, with larger LLRs and a decrease in lateral variability between peak velocity and movement end (0.73 to 0.63 cm), whereas variability increased with the wide target (1.11 to 1.94 cm). We also observed the same pattern, as well as the lower preparatory beta, in separate blocks without online visual feedback, suggesting reliance on proprioceptive and/or internal feedback rather than on vision alone. Furthermore, LLRs have been shown to increase with decision evidence (Selen et al., 2012) and reward (Codol et al., 2020; De Comite et al., 2022), conditions that also lower preparatory beta (Tzagarakis et al., 2010; Savoie et al., 2019). Such a link is also compatible with accounts in which beta reflects the weight afforded to sensory prediction errors (Palmer et al., 2016, 2019), as deviations become more consequential with a narrow target and should be weighted and corrected more strongly. This logic may also extend to preparation, as optimal control models propose that deviations in preparatory activity are themselves corrected by feedback, selectively along dimensions relevant to the upcoming movement (Kao et al., 2021). The quenching of preparatory variability described above could thus rely on internal feedback control. Lower beta could therefore accompany a greater allocation of resources to feedback control, attenuating both internal and external noise.

The topography of the effect is consistent with both possibilities. The condition difference was strongest over bilateral parieto-occipital electrodes, whereas the desynchronization common to both conditions was maximal over left centroparietal sites, suggesting that accuracy constraints modulated a posterior process superimposed on shared sensorimotor beta suppression. Posterior parietal cortex integrates goal- and effector-related information for upcoming movements (Andersen and Cui, 2009) and has been proposed to estimate the limb’s state from efference copies (Mulliken et al., 2008). Parietal area 5 also responds to perturbations before motor cortex when feedback is task-relevant (Omrani et al., 2016), and parietal disruption impairs online corrections (Desmurget et al., 1999; Takei et al., 2021). Although scalp EEG cannot identify generators, posterior circuits could thus contribute to both specifying and correcting the action.

### Alternative accounts

Reaction times were slightly shorter for narrow targets, and because lower preparatory beta is associated with readiness and faster responses (Kilavik et al., 2013), the effect could reflect a general state of readiness. However, the difference was small (13 ms), and readiness alone would not explain why movements were also more precise. Similarly, movement times were slightly longer for narrow targets, but neither peak velocity nor time to peak velocity differed, so the beta effect is unlikely to reflect a difference in planned movement speed. Finally, there was a rightward bias (2cm) in mean endpoint location with the wide target, a small difference that seems unlikely to account for the beta effect.

Alpha and theta showed the expected task-related dynamics, with posterior alpha desynchronization (Jensen and Mazaheri, 2010; Klimesch, 2012) and midfrontal theta increases (Cavanagh and Frank, 2014), but neither was reliably modulated by target width. Although null results should be interpreted cautiously, this suggests that higher precision demands were not expressed as a broad change in visuospatial attention or cognitive control.

### Limitations

Target width necessarily altered the visual display as well as accuracy constraints. Although a wide contour was overlaid beneath the narrow target, visual salience still differed. Three observations nonetheless argue against a purely visual explanation of the beta effect. First, condition-specific ERPs were removed before time-frequency decomposition, ruling out differences in phase-locked visual responses. Second, the beta difference emerged only in the second half of the delay, whereas differences driven by the display would be expected soon after target onset. Third, greater visuospatial attention to the narrow target should have produced stronger posterior alpha desynchronization (Thut et al., 2006), yet the largest, nonsignificant alpha cluster showed the opposite trend.

## Conclusion

Here, we show that with narrow targets, movements become more precise, through greater co-activation and larger LLRs, and preparatory beta power decreases. Preparatory beta is therefore sensitive to precision demands, and its decrease may accompany stronger quenching of preparatory variability, higher feedback gains, or both. Beyond where and how fast to move, movement preparation thus appears to take into account how precisely a movement must be executed.

## Acknowledgements

This work was supported by Fonds de recherche du Québec - Nature et technologies (FRQNT), grant 328238. The authors used Claude (Claude Opus 5.5; Anthropic) to improve the wording and clarity of the text. The authors reviewed and edited all suggested changes and take full responsibility for the content of the manuscript.

## Code availability

Custom MATLAB and Python analysis code is available upon reasonable request.

