## Extended data for "Preparatory beta activity reflects the precision demands of an upcoming movement"

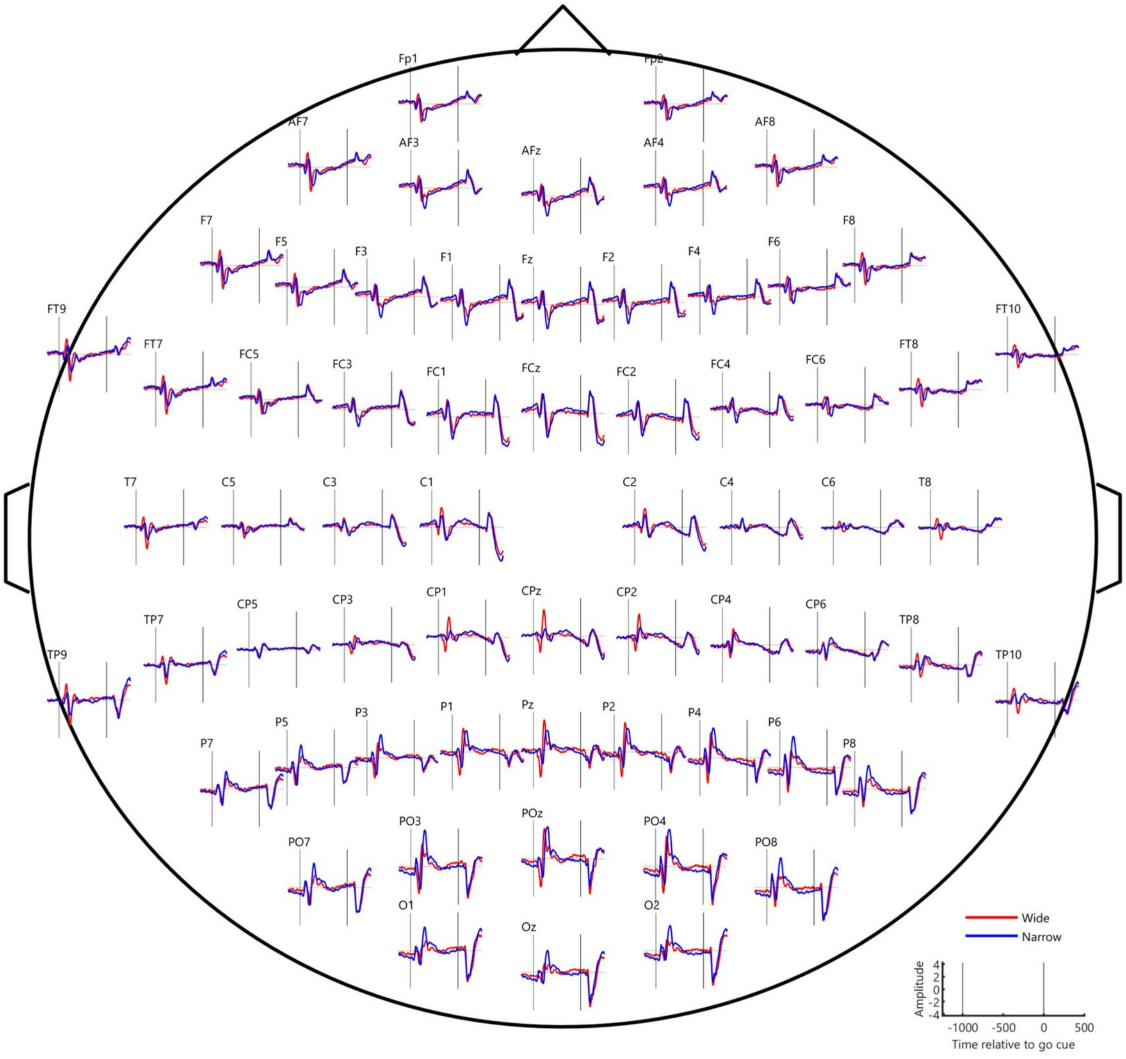


**Extended Data Figure 4-1. Event-related potentials following target presentation.** Grand-average ERPs at each electrode for the wide (red) and narrow (blue) targets, shown relative to the go cue. Electrodes are arranged according to their scalp position. n = 27.

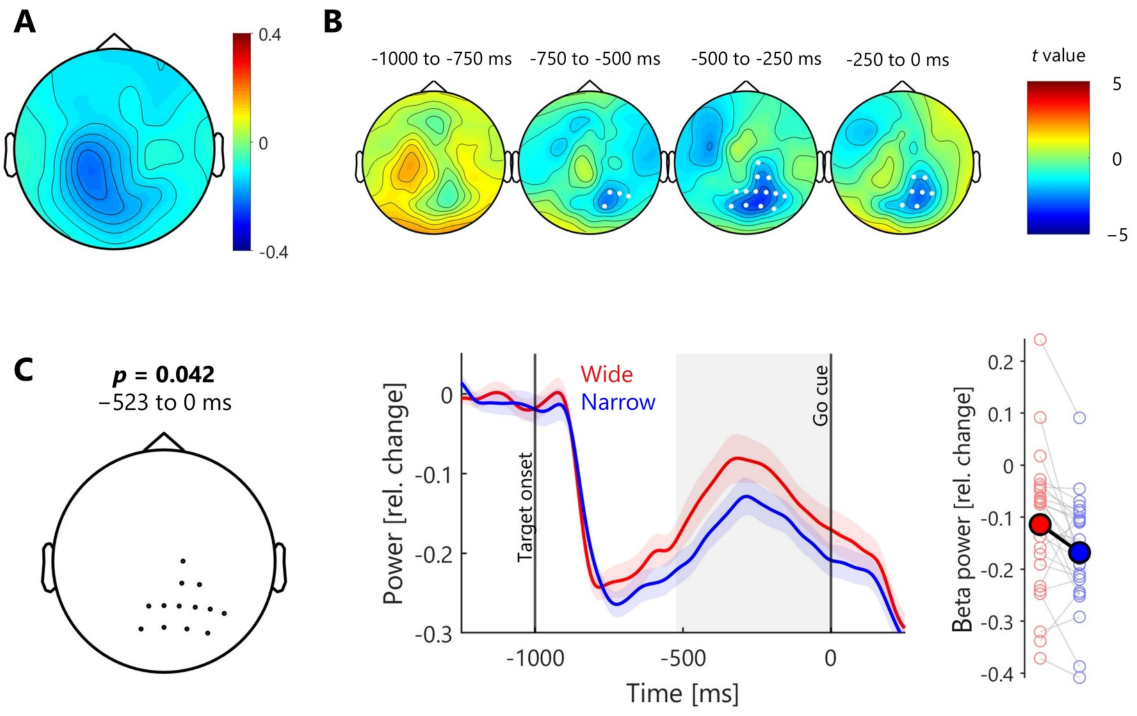


**Extended Data Figure 4-2. Preparatory beta power without visual feedback.** A, Topography of beta power (13-30 Hz, relative change from baseline) during the delay period, averaged across conditions. B, Topographies of the condition difference (narrow minus wide, t values) in four successive 250-ms windows preceding the go cue. White markers indicate electrodes belonging to the significant cluster. C, Left, electrodes of the significant cluster, with the cluster p value and time window. Middle, beta power averaged over cluster electrodes for the wide (red) and narrow (blue) targets, aligned to the go cue (mean ± SEM). The grey area indicates the time window of the cluster. Right, beta power averaged over cluster electrodes and time window for each participant (light markers) and the group mean (dark markers). Because electrodes were selected from the same contrast, the traces in C are descriptive (Kriegeskorte et al., 2009). n = 27.
